# Predictable Timing Enhances the Use of Predictable Content

**DOI:** 10.64898/2026.10.23.760967

**Authors:** Oscar Magnusson, Hanne Hobbelen, Wouter Berkhof, Alexander T. Sack, Inge Leunissen, Sanne Ten Oever

**Affiliations:** Department of Cognitive Neuroscience, Faculty of Psychology and Neuroscience, Maastricht University, Oxfordlaan 55, 6229 EV Maastricht, The Netherlands

**Keywords:** periodicity, autocorrelation, 1/f, auditory tracking, prediction

## Abstract

As events unfold, they carry structure in both their timing and content, with these dimensions often interrelated. Most research has treated these dimensions separately, although processing them jointly may provide important computational benefits. It is an open question whether the brain processes a sequence’s timing and predictable content structure independently, or integrates these two sources of information to guide prediction. We hypothesised that predictable timing would allow listeners to better exploit structure in the content, with this effect increasing as content structure becomes stronger. To test this, we presented tone sequences with pitch trajectories following a 1*/f* ^*β*^ structure, where *β* determines how strongly the content of each tone persists, while independently varying the temporal periodicity of the sequence. Performance was quantified using response errors relative to outcome predictions of a Gaussian process regression model and a model-free measure of how strongly preceding tones influenced responses. Response error decreased and reliance on preceding tones increased with higher *β* and periodic timing. Moreover, the influence of preceding tones showed an interaction between timing and content, such that the effect of periodicity grew with increasing *β*. The response error showed no such interaction, although the effect of periodic timing was reliable only for the most structured sequences. Our findings suggest an interdependency between content and timing, where predictable timing helps listeners make greater use of predictable content.

## Introduction

Natural sensory input carries structure in both its content and timing. The content of successive events can be related, such that some sequences stay stable and highly predictable while others change unpredictably. The timing of events can likewise vary, ranging from strictly periodic to fully irregular. Perception exploits both aspects of the structure, anticipating what is likely to occur and when it will arrive (Clark, 2013; de Lange et al., 2018).

Predictable timing sharpens sensory processing by prioritising moments when relevant input is expected (Nobre & van Ede, 2018). It can arise either from cue-target pairs with a predictable interval (Niemi & Näätänen, 1981) or from the extraction of regularity in rhythmic input (Large & Jones, 1999), with both improving perceptual sensitivity and response speed (Chang et al., 2019; Correa et al., 2005; Jones et al., 2002; Lawrance et al., 2014). Such alignment may increase neural gain for any stimulus arriving at the expected moment (Auksztulewicz et al., 2019), with temporal expectation shown to sharpen the precision of sensory representations at that moment (Rohenkohl et al., 2012).

Predictable content also improves perception. The brain readily extracts statistical regularities to anticipate upcoming input (Saffran et al., 1999; Turk-Browne et al., 2010), improving perceptual sensitivity (Cheadle et al., 2015; Wyart et al., 2012). Most paradigms have relied on strong, experimentally induced predictability, such as learned cue-target pairings (Summerfield & Egner, 2009) or repeated patterns in oddball paradigms (Näätänen et al., 2007). A more natural form of content predictability is autocorrelation, which captures how far a signal’s content persists over time. Speech, music, and environmental sounds show 1*/f* ^*β*^ power spectra (Gilden, 2001; Voss & Clarke, 1975), where higher values of *β* reflect stronger long-range correlations and thus higher predictability (Eke et al., 2000; He, 2014). Listeners exploit this structure (Garcia-Lazaro et al., 2006; Lin et al., 2016; Patel & Balaban, 2000; Schmuckler & Gilden, 1993) as predictions of upcoming tones draw on longer stretches of the preceding sequence when *β* is higher (Baumgarten et al., 2021; Maniscalco et al., 2018).

In natural input, what occurs and when it occurs are often bound together (Simoncelli & Olshausen, 2001), yet it remains unclear whether their predictability influences perception independently or interactively (ten Oever & Martin, 2024). Several findings point towards an interaction. Neural representations of predicted content peak at the moment that content is expected (Kaltenmaier et al., 2026), and predictable timing improves perceptual sensitivity particularly when the content itself is predictable (Morillon et al., 2016; Wetekam et al., 2026). However, this interaction may depend on task demands. Benjamin et al. (2026) found that temporal irregularity impaired the discovery of a latent rule in a visual prediction task but left sensory integration intact, suggesting that timing benefits only some forms of content-based prediction (see also Kim et al., 2025). This leaves two open questions. First, under which task demands do timing and content interact? Second, as most studies have treated content as either fully predictable or unpredictable, is this interaction graded, scaling with how strongly the content supports prediction?

Here, we investigate the interaction between content and temporal predictions. We presented listeners with pure-tone sequences whose pitch followed a 1*/f* ^*β*^ structure at three levels of *β*, under periodic or aperiodic presentation. We assessed both how closely listeners’ responses followed model-derived predictions and how much their responses were shaped by preceding tones. We hypothesised that the influence of temporal regularity would increase with *β*, being strongest when strong autocorrelation supports content prediction and weakest when content is less predictable.

## Methods

### Participants

Forty-one students from Maastricht University were recruited for the study (mean age = 20.49 ± 2.25 years, 9 males). All participants provided their informed written consent before participation. Participants reported no personal or family history of epilepsy, seizures, or convulsions, no neurological disorders, brain injury or stroke, no current or previous psychiatric disorders, and no diagnosed hearing impairments. The experimental protocol was approved by the ethical committee of the Faculty of Psychology and Neuroscience at Maastricht University (approval number OZL_264_26_02_2023) and was conducted in accordance with the Declaration of Helsinki. Participants received monetary compensation or course credit for their participation.

### Experimental stimuli

Auditory stimuli were presented as sequences of pure tones drawn from 25 discrete pitch levels spanning ±12 semitones around 440 Hz (220–880 Hz). Sequence length was randomly sampled on each trial (*n*_tones_ ∈ [20, 30]). The spectral structure of each sequence was manipulated at three levels of *β* (0.25, 0.99, and 1.75), where *β* controlled the slope of the 1*/f* ^*β*^ spectrum (Figure 1A). Increasing *β* shifted relatively more spectral energy towards lower frequencies, producing pitch trajectories with stronger autocorrelation and slower, more persistent pitches over time.

**Figure 1.**
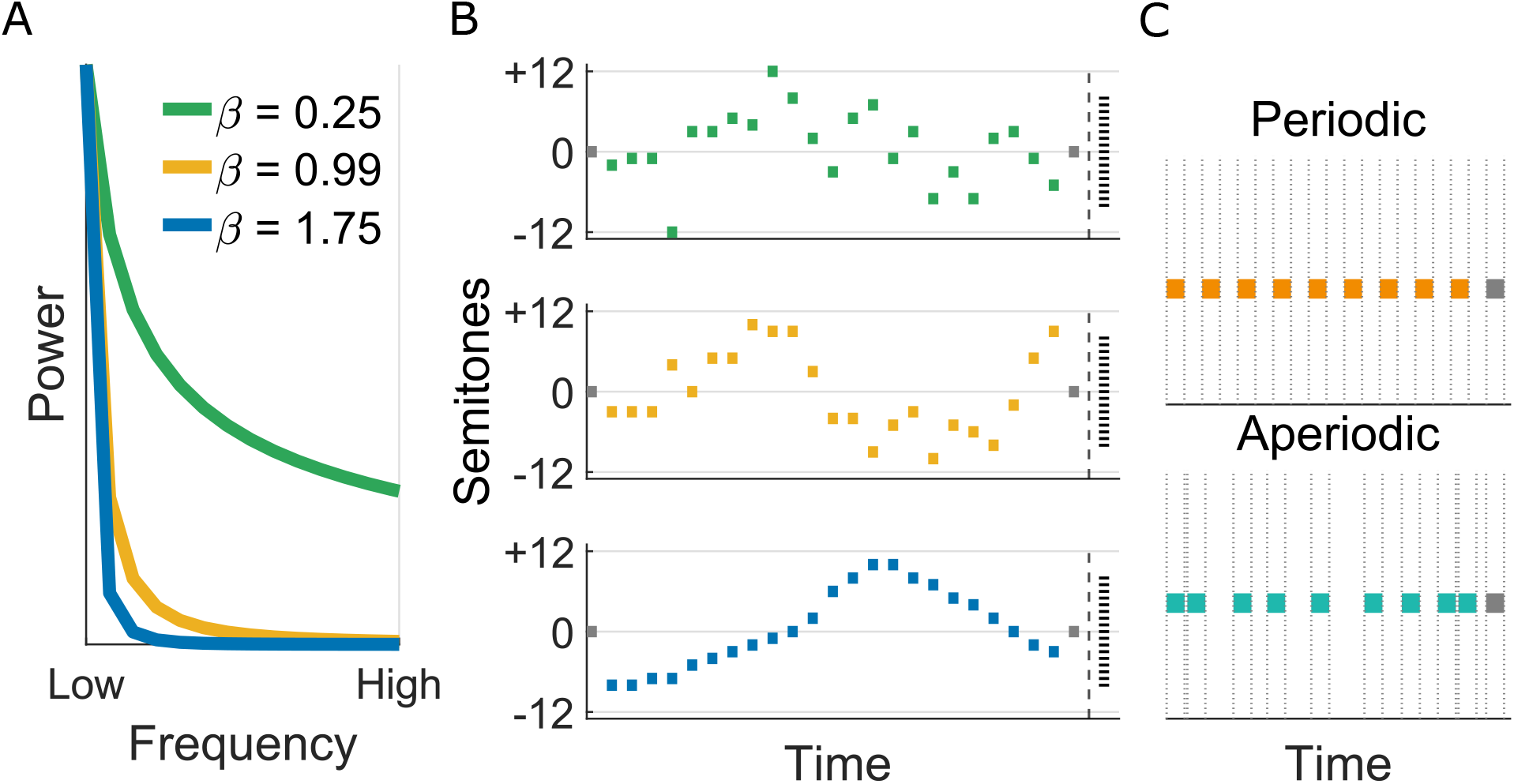
Illustration of the design. (A) Power spectra of tone sequences. Sequences followed a 1*/f*^*β*^ structure at one of three levels of the exponent *β*, where higher values shift spectral energy towards lower frequencies and give stronger autocorrelation and slower, more persistent pitch trajectories. (B) Pitch trajectories over time. Pitch spanned +12 to −12 semitones relative to 440 Hz, with grey anchor tones fixed at the start and end and coloured tones showing the structured trajectory. Participants responded on the scale from +8 to −8 semitones shown on the right. (C) Timing manipulation. Tones were always 150 ms, and in periodic sequences the inter-stimulus interval was fixed at 150 ms, whereas in aperiodic sequences it was jittered uniformly between 0 and 300 ms, keeping the same 150 ms mean rate.

The pitch trajectory for each trial was generated in the frequency domain. We first constructed the positive-frequency components of a discrete Fourier spectrum at a length *N* = 2*n*_notes_. We used twice the final sequence length, so that the frequency grid contained enough low-frequency components to represent the 1*/f* ^*β*^ structure faithfully, since at higher *β* almost all spectral power falls in the lowest frequencies. Fourier amplitudes were scaled according to the spectral exponent 1*/f* ^*β*^, and phases for the positive-frequency components were sampled independently from a uniform distribution over [0, 2*π*], so that each trial was a unique realisation of the same spectral process. The negative-frequency components were set as the complex conjugate mirror of the positive-frequency components, imposing Hermitian symmetry so that the inverse FFT returned a real-valued trajectory. After transforming to the time domain, we retained the first *n*_notes_ samples and discarded the remainder. This truncation serves two purposes. It removes the periodic boundary of the inverse FFT, where the end of the signal wraps onto its start, so that the retained segment is not forced to be cyclic, and it leaves a trajectory whose start and end can differ, as a genuine sequence should. The retained segment was then mean-centred and normalised to a standard deviation of 6 semitones, controlling the pitch range and minimising clipping at the boundaries, after which values were rounded to the nearest semitone (Figure 1B). Finally, the first and last tones were fixed at 0 semitones (440 Hz) to provide anchor references for the prediction task. Tone duration was fixed at 150 ms across all conditions. Inter-stimulus intervals (ISIs) were fixed at 150 ms in periodic sequences and drawn from a uniform distribution between 0 and 300 ms (mean 150 ms) in aperiodic sequences (Figure 1C). Each tone included a 10 ms linear amplitude ramp at onset and offset.

To estimate the expected next tone of each sequence, we fitted a Gaussian process regression (GPR) model separately to every trial (Rasmussen & Williams, 2006). GPR is a non-parametric Bayesian method that does not assume a fixed functional form, such as a linear trend, for the pitch trajectory. Instead, it places a prior distribution over possible functions relating tone position to pitch, defined by a mean function and a covariance function (kernel). The kernel specifies how strongly the pitches of two tones are expected to covary as a function of the distance between their positions, such that nearby tones are assumed to be more similar than distant ones. Conditioning this prior on the observed tones yields a posterior distribution over functions consistent with the sequence heard so far. Evaluated at a new position, this posterior provides a Gaussian predictive distribution for the upcoming tone. Its mean is a weighted combination of the observed tones, with weights determined by their kernel similarity to the predicted position, and its variance reflects the uncertainty of the prediction.

We used a rational quadratic kernel

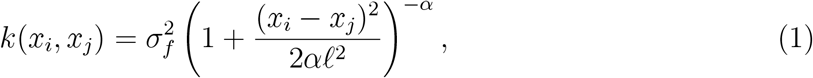

where *x*_*i*_ and *x*_*j*_ are tone positions, *σ*_*f*_ is the signal standard deviation, *l* is the characteristic length scale in tones, and *α* is a shape parameter. This kernel is equivalent to a mixture of squared exponential kernels with different length scales, with *α* determining the relative weighting of short and long scales (Rasmussen & Williams, 2006). It can therefore capture both slow global trends and faster local fluctuations, making it suitable for sequences ranging from weakly (*β* = 0.25) to strongly (*β* = 1.75) autocorrelated. The kernel parameters, together with a constant mean and the observation noise variance, were estimated separately for each trial by maximising the log marginal likelihood, so that the model adapted to the correlation structure of each individual sequence.

The model was trained on the intermediate tones of the sequence, excluding the anchor tones, with tone position as the predictor and pitch in semitones as the response. Pitch values were z-scored within each trial prior to fitting to improve numerical stability. Model fitting was performed using MATLAB’s (R2025b) `fitrgp` function (Statistics and Machine Learning Toolbox, version 12.4) with default settings (constant basis function, exact parameter estimation, and exact prediction). The fitted model was then evaluated at the position immediately following the last intermediate tone, and the mean of the resulting posterior predictive distribution was taken as the predicted pitch. This value was transformed back to semitone units and rounded to the nearest integer semitone, matching the resolution of the response scale.

Because sequences were generated stochastically, the target tone, defined as the GPR’s predicted next tone, emerged from each sequence rather than being set in advance. To balance the design, we generated sequences iteratively and retained only those whose predicted target matched the level required, until every combination was filled. The target range was restricted to ±8 semitones relative to 440 Hz (17 outcomes), as wider ranges could not be filled at all levels of spectral structure, extreme targets being rare at low *β*. Each combination of *β* (3 levels), periodicity (2 levels), and target semitone (17 levels) was presented five times, giving 510 trials per participant, in pseudo-randomised order.

### Paradigm and procedure

Participants first completed 34 training trials using highly structured periodic sequences (*β* = 1.75, fixed ISI of 150 ms), covering two repetitions of each semitone outcome from −8 to +8. Feedback showing both the selected and correct semitone was provided after each training trial. In the main experiment, sequences were presented while a white fixation cross was displayed on a grey screen. After each sequence, a 1 s blank screen preceded the response phase, in which participants reported their predicted next tone on a vertical semitone scale ranging from −8 to +8 (17 discrete options) using a mouse, followed by a confidence rating on a horizontal scale from 0 to 100. Trials were self-paced. The 510 trials were divided into 15 blocks of 34 trials each, split across two sessions on separate days (approximately 1.5 hours per session).

The experiment was presented using MATLAB (R2024b, 24.2.0.2712019) and Psychtoolbox (version 3.0.19) and conducted in a soundproof room. Visual stimuli were presented on an Iiyama ProLite XUB2493HSU monitor (1920 × 1080 pixels, 60 Hz) at an approximate viewing distance of 80 cm without headrest. Auditory stimuli were delivered diotically via Sennheiser HD650 headphones connected to a Focusrite Forte USB audio interface, at a sampling rate of 48 kHz and 24-bit resolution. Stimuli were presented at a fixed level of approximately 65 dB SPL, identical for all participants.

## Data analysis

### Reference model validations

To validate GPR as a reference for correct responses in participants’ prediction errors, we compared trial-wise GPR output against a simple linear baseline, whose outputs were likewise rounded to the nearest semitone. The linear model captures the overall linear trend of the sequence, while GPR additionally models local and non-linear structure. We first compared the two models by correlating their outputs with the responses of the participants, pooling all trials. We then tested the difference between these correlations with a Steiger’s *Z*-test (Steiger, 1980). Next, we computed for each trial the absolute deviation of each model’s output from the participant’s response and calculated the deviation difference of the linear and GPR model such that a positive value indicates the GPR output fell closer to the response than the linear output. We tested the average advantage of the GPR model with an intercept-only linear mixed-effects model with the difference as dependent variable and including a random intercept for participant. Additionally, we added *β*, periodicity, and their interaction as fixed effects to test for variation of this advantage across content and time structure conditions. Simple effects at each level of *β* and periodicity were estimated with marginal means.

### Response error and confidence

Response error was defined as the trial-by-trial absolute difference in semitones between the GPR-predicted tone and the participant’s response. Response errors and confidence ratings were analysed in R (version 4.4.2) using generalised and linear mixed-effects models, respectively. Response errors were modelled using a negative binomial generalised linear mixed-effects model with a log link, appropriate for right-skewed, overdispersed data such as the absolute response errors on the semitone scale. Confidence ratings were modelled using a standard linear mixed-effects model based on a Gaussian distribution. Both models included *β* (0.25, 0.99, 1.75) and periodicity (periodic, aperiodic) as fixed effects, with *β* = 0.25 aperiodic as reference level. Both models included by-participant random intercepts. A by-participant random slope for *β* was considered but not retained, as it produced singular fits and was therefore dropped in favour of a random-intercept structure. For confidence, effect sizes are reported using 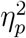. For response error, effect sizes are reported as rate ratios with 95% confidence intervals as no 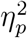 can be calculated from a negative binomial model. Hence, the effect sizes between these models are not comparable.

### Pitch tracking

In the response error analysis, performance was assessed relative to the output of the GPR model. However, sequences were generated stochastically from a 1*/f* ^*β*^ process, so no tone can be designated as the objectively correct continuation on a given trial, and responses cannot be scored for accuracy against a known target. We therefore performed a complementary analysis that relates responses directly to the sequence itself. The resulting pitch–response slope is a model-free index of how strongly the preceding tone influenced participants’ responses. A slope of 1 indicates that a one-semitone change in the preceding tone shifted the response by a full semitone, and a slope of 0 indicates no influence.

Responses were modelled at the trial level with a linear mixed-effects model in lme4 (Bates et al., 2015), specified as

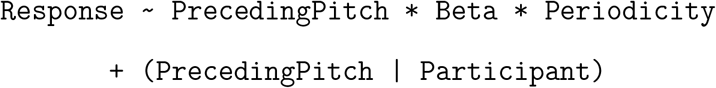

Written out, the response *R*_*ij*_ on trial *i* of participant *j* is

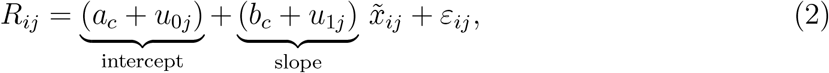

where 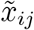 is the mean-centred pitch of the immediately preceding tone (the last non-anchor tone before the response) and *c* denotes the *β×* `Periodicity` condition of the trial. The model thus fits one line relating preceding pitch to response per condition, and conditions can differ in two ways. The slope *b*_*c*_ reflects pitch tracking, i.e., how strongly responses follow the preceding tone. The intercept *a*_*c*_ reflects baseline responding, i.e., whether responses are shifted up or down overall, regardless of the preceding tone. The terms *u*_0*j*_ and *u*_1*j*_ allow each participant their own intercept and slope, and *ε*_*ij*_ is the residual error. Mean-centring preceding pitch places the intercept at the average preceding pitch rather than at an arbitrary 0 semitones, so that the two components can be interpreted separately. In the model formula, effects involving `PrecedingPitch` therefore describe the slope, and the remaining condition effects describe the intercept. Our central question, whether content predictability modulates the effect of periodicity on pitch tracking, is thus tested by the `PrecedingPitch` *× β ×* `Periodicity` interaction.

To determine whether a by-participant random slope for preceding pitch was warranted, we compared the model against an otherwise identical model with random intercepts only, using a likelihood ratio test. All reported effects were obtained from the final model. Fixed effects were tested with Type III *F* -tests using Satterthwaite degrees of freedom (Kuznetsova et al., 2017), with sum-coded factors. Condition-specific slopes and contrasts were obtained with `emtrends` and `emmeans` (Lenth & Piaskowski, 2025), with Holm correction, and periodicity contrasts are reported as periodic minus aperiodic.

Unlike serial dependence studies using independently sampled stimuli, we did not analyse deeper lags as an index of integration, because the generative process makes successive tones strongly autocorrelated and collinear, increasingly so at higher *β*, with adjacent-tone correlations of .36, .83 and .97 at *β* = 0.25, 0.99 and 1.75. Entering multiple lags jointly was therefore untenable at high *β*, where variance inflation factors reached 117.8 and the deeper tones carried almost no unique information, with no lag beyond the second adding more than 1.3 % of response variance in any condition. None of this applies to the most recent tone, which is estimated in a model containing no other lag, and autocorrelation was closely matched between periodic and aperiodic sequences within each level of *β*, differing by at most Δ*r* = .02, so the periodicity contrast within a level of *β* is unconfounded by sequence structure.

## Results

### Reference model validations

GPR predictions were shown to correlate more strongly with participants’ responses (*r* = .566, 95% CI [.557, .576], *p <* .001) as compared to the linear baseline model (*r* = .515, 95% CI [.505, .525], *p <* .001; Steiger’s *Z* = 9.16, *p <* .001; see Figure 2). The linear mixed-effect model confirmed that the GPR output was, on average, 1.76 semitones closer to participants’ responses than the linear model (*b* = 1.76, *SE* = 0.07, *t*(40) = 23.62, *p <* .001). This advantage was modulated by *β* (*F* (2, 20864) = 1638.38, *p <* .001), being negligible at *β* = 0.25 (0.05 semitones) but increasing at *β* = 0.99 (1.87) and *β* = 1.75 (3.35). Surprisingly, it was slightly larger for aperiodic than periodic sequences, with a 0.11 semitone improvement (*F* (1, 20864) = 5.09, *p* = .024). No *β ×* periodicity interaction effect was observed (*F* (2, 20864) = 1.86, *p* = .155). GPR thus provides a reference closer to participants’ responses than the linear baseline model, particularly as *β* increases.

**Figure 2.**
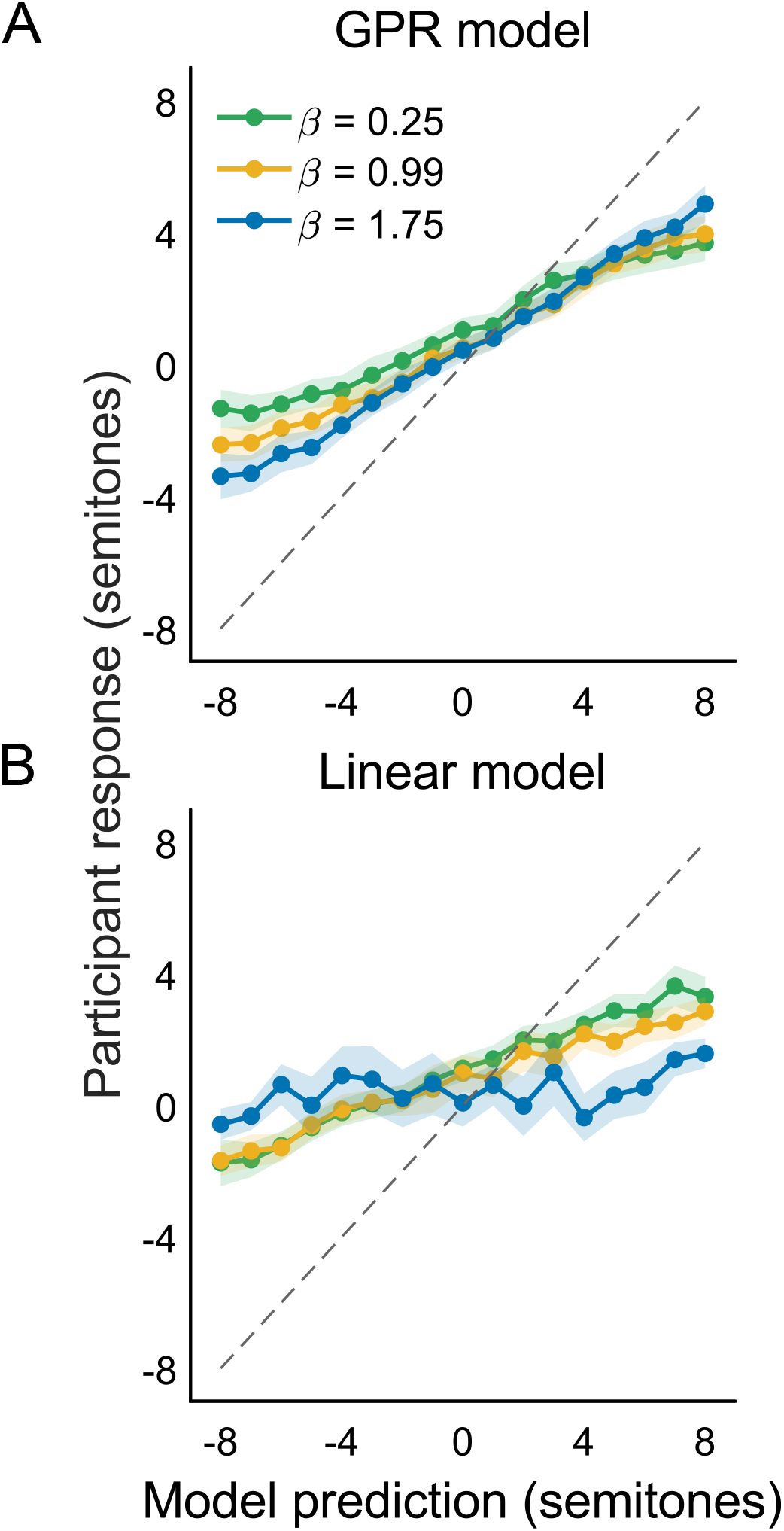
Mean responses as a function of model-predicted semitone across *β* levels, for the GPR model (A) and the linear model (B). Points indicate means with shaded regions giving the ±95% CI across participants. The dashed diagonal marks perfect correspondence between model predicted next tone and response.

### Response error

We defined response error as the absolute difference on each trial between a participant’s response and the GPR-estimated semitones, and asked whether it varied with condition. The response error was shown to decrease with higher *β* (*χ*^2^(2) = 253.61, *p <* .001), where the errors were 0.72 semitones lower for *β* = 1.75 than *β* = 0.25 (95% CI [0.51, 0.92], *p <* .001). They were 0.41 semitones lower for *β* = 0.99 than for *β* = 0.25 (95% CI [0.24, 0.57], *p <* .001) and 0.31 semitones lower for *β* = 1.75 than *β* = 0.99 (95% CI [0.20, 0.42], *p <* .001; see Figure 3A). Response errors were also 0.11 semitones lower for periodic than aperiodic sequences (95% CI [0.04, 0.18], *χ*^2^(1) = 9.19, *p* = .002).

**Figure 3.**
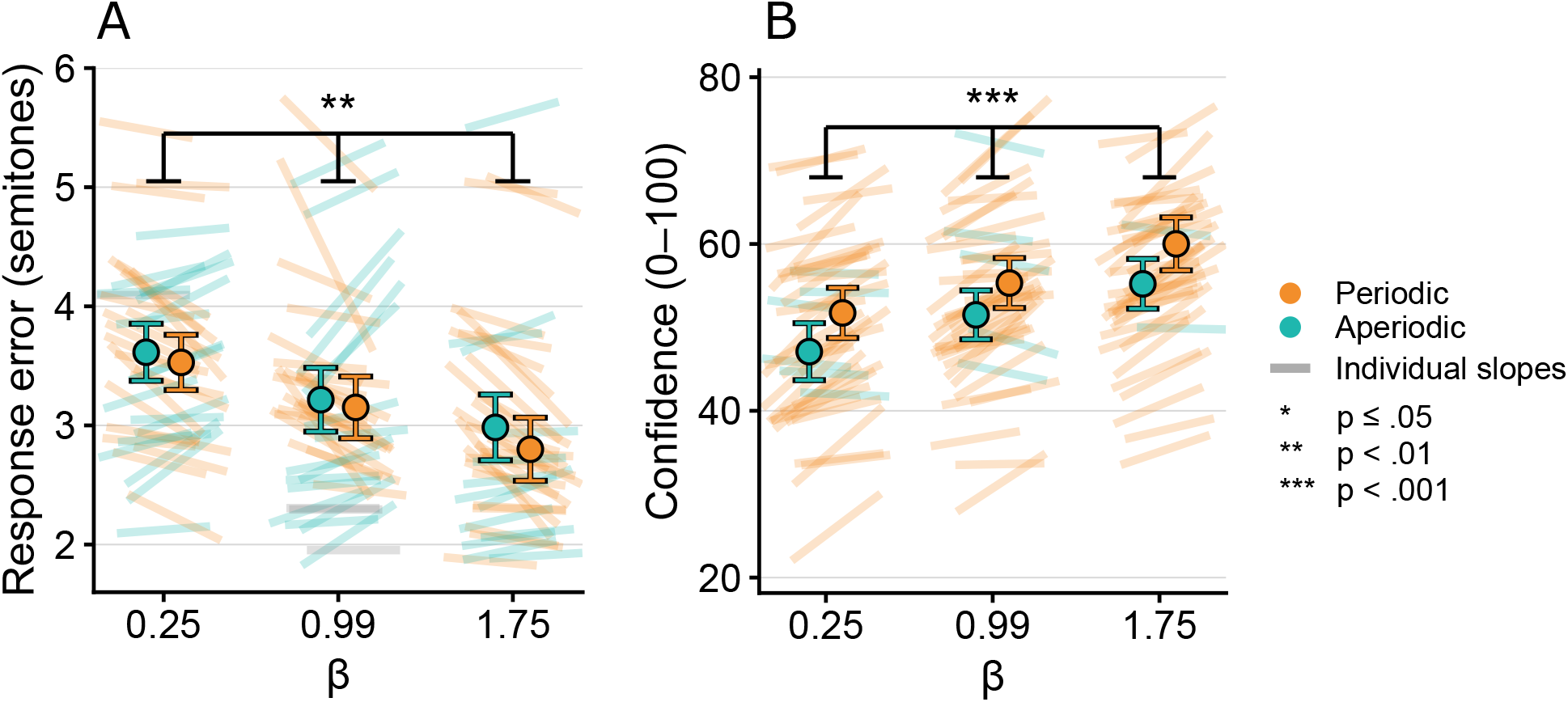
Response error and confidence ratings across *β* levels and periodicity condition. Points indicate condition means (± 95% CI). Lines connect individual participants’ values across aperiodic and periodic conditions within each *β* level. (A) Response error: orange lines indicate lower response error in the periodic relative to the aperiodic condition (downward slopes), teal lines indicate the opposite pattern (upward slopes). (B) Confidence ratings: orange lines indicate higher confidence in the periodic relative to the aperiodic condition (upward slopes), teal lines indicate the opposite pattern (downward slopes).

As planned, we tested whether the periodicity benefit depended on the sequence content structure set by *β*. The *β ×* periodicity interaction was not significant (*χ*^2^(2) = 2.81, *p* = .245), indicating that the periodicity effect did not differ reliably across the levels of *β*. Planned within-*β* contrasts showed a reliable benefit only at *β* = 1.75, where the response error was 0.18 semitones lower under periodic timing (95% CI [0.06, 0.29], *p* = .002), with smaller, non-significant differences at *β* = 0.25 (0.09 semitones, 95% CI [−0.05, 0.22], *p* = .203) and *β* = 0.99 (0.06 semitones, 95% CI [−0.06, 0.18], *p* = .348).

### Confidence ratings

Confidence ratings, given after each trial on a scale from 0 to 100, were also compared across conditions. Participants’ confidence increased with higher *β* (*F* (2, 20869) = 357.47, *p <* .001, 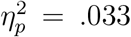), having 8.20 higher confidence points for *β* = 1.75 than *β* = 0.25 (95% CI [5.60, 10.81], *p <* .001, 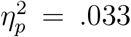). Their confidence was 4.00 points higher for *β* = 0.99 than *β* = 0.25 (95% CI [2.29, 5.70], *p <* .001, 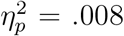), and 4.21 points higher for *β* = 1.75 than *β* = 0.99 (95% CI [2.95, 5.47], *p <* .001, 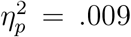; see Figure 3B). Confidence was also 4.42 points higher for periodic than aperiodic sequences (95% CI [3.93, 4.90], *F* (1, 20869) = 310.97, *p <* .001, 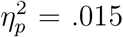). The *β ×* periodicity interaction was not significant (*F* (2, 20869) = 1.46, *p* = .232).

### Pitch tracking

To assess how strongly listeners carried the preceding tone into their responses, we estimated pitch–response slopes from a trial-level model of responses on the semitone of the immediately preceding tone. Allowing pitch tracking to vary across participants substantially improved model fit (*χ*^2^(2) = 1914.6, *p <* .001), indicating reliable individual differences in how strongly listeners followed the preceding tone, and the random slope was therefore retained. We report slope-level effects, reflecting pitch tracking, followed by intercept-level effects, reflecting baseline responding.

Listeners tracked the preceding tone reliably to inform their final response (*F* (1, 41) = 294.94, *p <* .001, 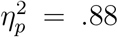). This tracking increased with *β* (*F* (2, 20829) = 871.50, *p <* .001, 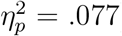). Comparing the model-estimated slopes across levels of *β*, the slope at *β* = 1.75 was 0.32 higher than at *β* = 0.25 (95% CI [0.30, 0.34], *p <* .001, 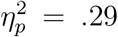), so a one-semitone shift in the preceding tone was carried 0.32 semitones further into the response as the sequence became more structured. This increase was graded, the slope being 0.15 higher at *β* = 0.99 than *β* = 0.25 (95% CI [0.13, 0.17], *p <* .001, 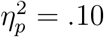) and a further 0.17 higher at *β* = 1.75 than *β* = 0.99 (95% CI [0.15, 0.20], *p <* .001, 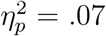; see Figure 4). Listeners thus weighted the most recent tone more heavily as the predictive nature of the sequence grew. Tracking was also stronger for periodic than aperiodic sequences (*F* (1, 20829) = 86.11, *p <* .001, 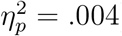), the model-estimated slope being 0.06 higher under periodic timing (95% CI [0.048, 0.073], *p <* .001), so the preceding tone was carried 0.06 semitones further into the response when timing was regular.

**Figure 4.**
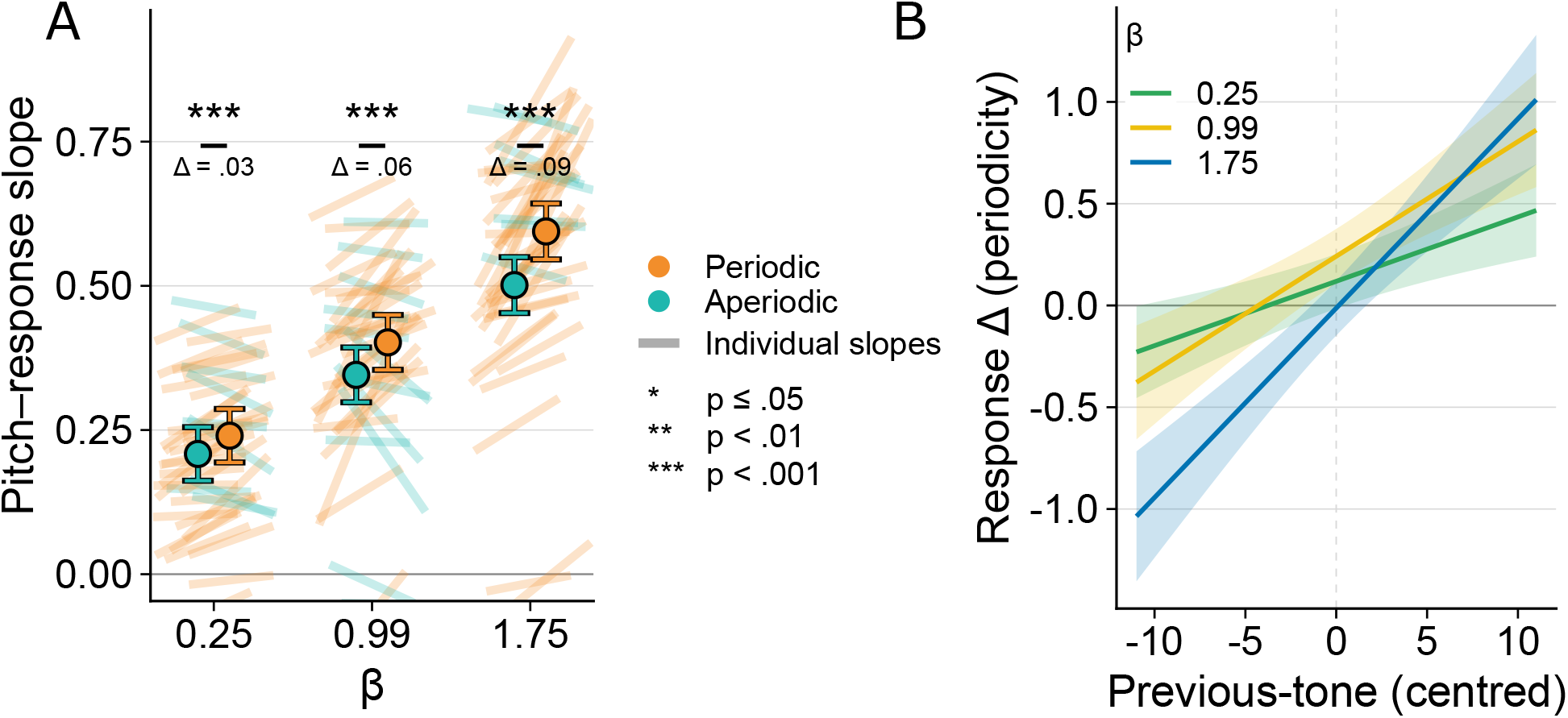
Pitch tracking results. A) Pitch-response slopes for the preceding tone, by spectral structure and periodicity. The slope indexes the degree to which a one-semitone change in the immediately preceding tone shifts a participant’s prediction of the next tone, shown for each *β* level and periodicity condition. Points give condition means (± 95% CI). Lines connect individual participants’ slopes between periodicity conditions within each *β* level, orange indicating stronger pitch tracking (a higher slope) in the periodic relative to the aperiodic condition (upward lines) and teal indicating the opposite pattern (downward lines). B) Difference in predicted response between periodic and aperiodic sequences (periodic minus aperiodic) as a function of the centred preceding-tone pitch, shown separately for each *β* level (*β* = 0.25 green, *β* = 0.99 yellow, *β* = 1.75 blue). Its height at zero reflects the periodicity effect on baseline responding (the intercept) and its tilt reflects the periodicity effect on pitch tracking (the slope; ± 95% CI).

Critically, there was an interaction between periodicity and *β* (*F* (2, 20829) = 7.67, *p <* .001, 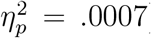). We found that the model-estimated periodicity influence across levels of *β* increased monotonically from 0.032 at *β* = 0.25 (95% CI [0.015, 0.048], *p <* .001, 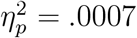) to 0.056 at *β* = 0.99 (95% CI [0.034, 0.079], *p <* .001, 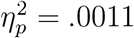) and 0.093 at *β* = 1.75 (95% CI [0.067, 0.119], *p <* .001, 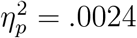). As content predictability increased, periodic timing strengthened the extent to which changes in the preceding tone were reflected in listeners’ responses.

This modulation acted on tracking rather than on baseline responding. Whereas the slope-level interaction was reliable, the corresponding intercept-level interaction was smaller and did not follow a graded pattern across *β* (*F* (2, 20828) = 3.45, *p* = .032, 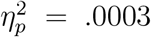). Comparing the model-estimated periodicity shift in baseline responding across levels of *β*, it was present at *β* = 0.99 (0.24, 95% CI [0.11, 0.38], *p <* .001) but not at *β* = 0.25 (0.12, 95% CI [−0.02, 0.25], *p* = .083) or *β* = 1.75 (−0.01, 95% CI [−0.15, 0.12], *p* = .849). Thus, periodic timing increased the influence of the most recent tone on listeners’ responses in proportion to content predictability, whereas its effect on baseline responding was smaller and showed no such graded pattern. The content-timing interdependence was therefore specific to pitch tracking rather than reflecting a general shift in responding.

## Discussion

We investigated whether listeners’ prediction of an upcoming tone’s pitch was shaped by the periodicity of tone presentation and by content predictability, manipulated through the sequence’s 1*/f* ^*β*^ structure, and whether the influence of periodicity increased with content predictability. Participants’ responses were more consistent with a predictive model of tone continuation for higher *β* and for periodic compared with aperiodic presentation. A model-free analysis of how strongly participants weighted the preceding tone in their responses showed corresponding main effects, with greater weight at higher *β* and under periodic presentation. Importantly, the two factors interacted, with the effect of periodicity on tracking growing monotonically with *β*. The model-based response error analysis showed no such interaction, although its periodicity benefit was reliable only at the most structured level (*β* = 1.75), a pattern broadly consistent with the tracking result.

Our findings add to a growing body of work showing that the predictability of content and timing shapes perception and prediction interdependently (ten Oever & Martin, 2024). Timing has been shown to carry content-specific predictions, both behaviourally, serving as a cue to stimulus identity (ten Oever & Sack, 2019; van de Ven et al., 2017), and neurally, with the representation of predicted content peaking when each item is due (Kaltenmaier et al., 2026). These effects, however, relied on learned associations between particular contents and particular moments. Our sequences contained no such associations, yet content and timing still interacted in shaping predictions. Our findings therefore align more closely with work showing that jointly predictable content and timing improve perceptual sensitivity beyond the sum of their individual effects (Morillon et al., 2016; Wetekam et al., 2026; Wollman & Morillon, 2018). We extend this work in two ways. First, the interaction emerged in a task requiring listeners to track naturally evolving structure, here the autocorrelation of 1*/f*^*β*^ sequences, identifying this as one demand under which timing and content interact. Second, the interaction was graded rather than arising from an all-or-nothing contrast between predictable and unpredictable input.

Not every task demand yields such a benefit, however. In a recent visual prediction task, in which participants predicted the orientation of the next Gabor grating in a stream, disrupting periodicity left sensory integration unaffected (Benjamin et al., 2026). Grating orientations were drawn from a distribution with a fixed mean, except when an unsignalled flip of the mean occurred partway through a sequence, a latent rule that only some participants discovered. Disrupting periodicity impaired the discovery of this flip while leaving integration of the mean intact. This suggests that temporal regularity aids processing when individual items are informative about upcoming input. When integrating the mean, each grating is an independent draw and contributes only marginally to the estimate, so precise processing of any single item adds little. At the flip, by contrast, a single item signals a change in all items to come, so precise processing at the expected moment carries a large predictive benefit. Our sequences fall between these extremes. Because of their autocorrelation, every tone is informative of the next, and raising *β* narrows the range in which tone *x* falls given tone *x* − 1. Precise processing of each tone at its expected moment should therefore yield a better prediction of the tone to come, with a benefit that grows with how strongly each tone constrains the next. We propose that this is what the graded content-timing interdependence reflects, with predictable timing being most beneficial when each item carries the most information about what follows.

An ideal observer should, however, draw not only on the most recent event but also on preceding input. Empirically, these events typically show a graded recency effect, with the immediately preceding event weighted most strongly and weights decreasing with increasing temporal distance into the past (Fischer & Whitney, 2014). Neural activity tracks just this for 1*/f* sequences, integrating up to seven preceding tones with a weighting profile matching the optimal weights set by the sequence’s autocorrelation (Maniscalco et al., 2018), an integration that depends on the number of preceding events rather than the time span they cover (Baumgarten et al., 2021). We would expect the content-timing interdependence to follow the same graded weighting of preceding events, with predictable timing enhancing the use of each event in proportion to its predictive value. Unfortunately, we cannot recover that graded decay from our responses in the current study design as the strong autocorrelation makes each tone highly predictive of the next, causing too strong collinearity to run our models.

There has been a long debate about whether many apparent prediction effects are more easily explained by lower-level adaptation than by top-down prediction (Grotheer & Kovács, 2016; Todorovic & de Lange, 2012). This could hold for our autocorrelated sequences. Rather than reflecting genuine prediction, stronger responses to structured than to unstructured sequences could arise because similar tones recur more often in structured sequences, repeatedly driving the same neural populations and inducing stronger adaptation (Nelken & Moshitch, 2026). The same logic could extend to timing, where regularly recurring arrivals adapt responses to that moment. Adaptation, however, predicts the opposite of what we observe. It weakens a response as content or timing repeats, so more structured and more regular sequences should yield weaker processing and lighter weighting of recent tones. We find the reverse, with lower response error and stronger tracking as content and timing become more predictable. Adaptation therefore cannot account for the direction of our effects, let alone their graded interaction.

While low-level adaptation is unlikely to explain our effects, other early-sensory mechanisms could potentially drive our content-timing interaction. Tone sequences in the delta-to-theta range, like ours, have been shown to entrain low-frequency oscillations with improved perceptual performance (Cravo et al., 2013; Herbst & Obleser, 2019; Lakatos et al., 2008; Stefanics et al., 2010; ten Oever et al., 2017). Such effects may arise, at least in part, from bottom-up sensory-driven mechanisms (Haegens & Zion Golumbic, 2018). The auditory cortex has a well-defined tonotopic organisation, where neuronal populations are selectively tuned to preferred auditory frequencies (Merzenich & Brugge, 1973) and mapped onto the cortical surface, with frequencies close in spectrum represented closer together in cortex (Da Costa et al., 2011; Humphries et al., 2010). This matters for our sequences, where higher *β* raises not only structuredness through autocorrelation but also the spectral proximity between successive tones, so tone *x* falls closer in frequency to tone *x* − 1. On the tonotopic map, this places the populations representing successive tones closer together. If entrainment to the rhythm engages one such population, activation of tone *x* − 1 could propagate to the neighbouring population representing *x*, so that the benefit of entrainment for the next tone grows as the sequence becomes more spectrally structured. Such propagation across topography has long been discussed, with evidence for spread across the retinotopic map (Cruddas et al., 2026; Petras et al., 2025; Sato et al., 2012). Evidence for tonotopic propagation is more limited. Horizontal projections between cortical populations exist (Kaur et al., 2005), and waves crossing the tonotopic axis have been observed after broadband and cochlear implant stimulation (Kral et al., 2009; Reimer et al., 2011), but tone-evoked propagation is mostly reported within isofrequency bands (Song et al., 2006; Versnel et al., 2002). Local propagation within auditory cortex could therefore contribute to the content-timing interaction, provided that activity evoked by one tone spreads to populations representing neighbouring tones.

An alternative, though not mutually exclusive, explanation is that top-down processes modulate sensory entrainment according to the predictability of the sequence content. In particular, entrainment can be shaped by top-down signals: attention has been shown to align the high-excitability phase of neural oscillation with an attended auditory stream, while suppressing competing streams at the low-excitability phase (Barczak et al., 2018; Lakatos et al., 2013). Although our current paradigm does not involve competing sensory input streams, similar top-down mechanisms could potentially modulate entrainment as a function of content predictability. The interaction shown here could therefore reflect a combination of entrainment to the stimulus rhythm and predictions about upcoming content. Entrainment may enhance processing at the expected moment of each tone (Auksztulewicz et al., 2019), while a predictive model of the unfolding sequence may improve sensitivity to predicted content (Bianco et al., 2026; Cope et al., 2017). As *β* rises and successive pitches become more constrained, these predictions would become more precise, increasing the benefit of processing each tone at its expected moment.

In conclusion, our results show that predictable timing increases the weight listeners assign to the most recent tone, with this effect becoming stronger as the sequence’s content becomes more predictable. This interaction suggests that content and temporal predictability jointly shape perception, rather than exerting independent, additive effects. Because auto-correlation makes preceding tones more informative about upcoming ones, regular timing allows listeners to make greater use of this predictive information. Together, these findings highlight how perception draws on the joint structure about what occurs and when it occurs to anticipate upcoming sensory events.

## Acknowledgements

This work was funded by a European Research Council grant (101116685) and a Dutch Research Council (NWO) grant awarded to STO. During the preparation of this manuscript, the authors used Grammarly and Claude (Anthropic) for language editing, and take full responsibility for the final text.

